# RNA-based vaccine demonstrates prophylactic efficacy against *Mycobacterium tuberculosis* challenge in a mouse model

**DOI:** 10.1101/2022.02.23.481669

**Authors:** Sasha E. Larsen, Jesse H. Erasmus, Valerie A. Reese, Tiffany Pecor, Jacob Archer, Amit Kandahar, Fan-Chi Hsu, Steven G. Reed, Susan L. Baldwin, Rhea N. Coler

## Abstract

*Mycobacterium tuberculosis* (Mtb) is an opportunistic bacterial pathogen that causes tuberculosis disease (TB) and exerts an extensive burden on global health. The complex intra- and extracellular nature of this bacterium, coupled with different disease stages have made mechanistic studies evaluating the contributions of innate and adaptive host immunity challenging. In this work we leveraged two delivery platforms as prophylactic vaccines to assess immunity and subsequent efficacy against low dose and ultra-low dose aerosol challenge with Mtb H37Rv in C57BL/6 mice. Our proof-of-concept TB vaccine candidate ID91 was produced as a fusion protein formulated with a synthetic TLR4 agonist (glucopyranosyl lipid adjuvant in a stable emulsion) or as a replicating-RNA (repRNA) formulated in a nanostructured lipid carrier (NLC). Results from this work demonstrate that protein subunit- and RNA-based vaccines preferentially elicit cellular immune responses to different ID91 epitopes. In a single prophylactic immunization screen, both platforms reduced pulmonary bacterial burden compared to controls. Excitingly, in prime-boost strategies, groups that received heterologous RNA-prime, protein-boost or combination (simultaneous in different sites) immunizations demonstrated the greatest reduction in bacterial burden and a unique humoral and cellular immune response profile. These data are the first to report that repRNA platforms are a viable system for TB vaccines and should be pursued with high priority Mtb antigens containing CD4+ and CD8+ T cell epitopes.

## Introduction

For the first time in a decade, 2020 saw an increase in annual deaths (1.5 million total) caused by *Mycobacterium tuberculosis* (Mtb)^1-3^. Before the Coronavirus disease 2019 (COVID-19) pandemic, caused by severe acute respiratory syndrome coronavirus 2 (SARS-CoV-2)^4^, Mtb was the world’s top cause of death from an infectious disease^5^. Exposure to Mtb can result in productive infection and pulmonary tuberculosis disease (TB). COVID-19 pandemic related disruptions in TB care and case finding are estimated by the World Health Organization (WHO) to lead to a further half-million preventable TB deaths^6^. This dual assault from respiratory pathogens is most heavily felt by low and middle income countries (LMICs) which endure a disproportionate burden of TB disease^7^, and are also suffering from a lack of access to vaccines for SARS-CoV-2^8^. Moreover, within Mtb, drug resistance (DR) is steadily increasing globally. For the past five years, nearly 0.5 million patients infected with Mtb annually develop resistance to front-line antibiotic rifampicin and approximately 80% of those harbor multidrug-resistance (MDR)^1,2,5^. Novel antibiotic regimens^9-13^ are being developed but by many models are insufficient alone to combat this epidemic^14,15^. Specifically designed low cost and effective TB vaccines for prevention of infection (POI), or complementary prevention of active disease (POD) endpoints are desperately needed^16^.

While clinical trials suggest that *Mycobacterium bovis* bacille Calmette-Guérin (BCG) may prevent Mtb infection in adolescents^17^, and subunit adjuvanted vaccine candidate M72/AS01_E_^18,19^ prevents TB disease in a subset of interferon gamma release assay (IGRA) positive individuals, there is no currently approved vaccine for POI or POD in adults. This lag is partially due to the fact that financial resources available for testing drug and vaccine candidate regimens in human clinical settings are disproportionately low for TB, relative to the global burden of disease. Addressing improved efficacy of prophylactic vaccines against Mtb and supply and costs of manufacturing materials are all required to ensure that a vaccine is readily available to those who need it. Thankfully, renewed enthusiasm for TB vaccine research has been stimulated largely by two main pillars of preclinical and clinical advancements. First, the pursuit of correlates of protection (COP) against TB which are refining the immune players in the fight against infection and disease, and second, the pursuit of cutting edge vaccine delivery platforms.

Historically the vaccine pipeline has relied on CD4+ T helper 1 (TH1) type responses as a benchmark for immunogenicity stage-gating of vaccine candidates. However, the full mechanism of protection has yet to be determined^20^, and several preclinical and clinical reviews suggest the CD4+ TH1 subset producing proinflammatory cytokine IFNγ alone is not sufficient nor fully predictive of clinical efficacy^21-24^. Indeed, protection from Mtb infection and TB disease is likely a multifaceted process involving many cell types beyond canonical CD4+ T cells, and our focus here is on Mtb-antigen specific CD8+ T cells. Many reviews are devoted to the significance of CD8+ T cells in TB infection and disease^25-30^, yet still they are not prioritized as a vaccine target cell type in most candidate screens. Importantly, CD8+ T cells can migrate to the site of Mtb infection^31-33^, and removing MHC class I or CD8+ T cells *in vivo* enhances Mtb susceptibility^28,34-37^. Cytolytic CD8+ T cells produce IFNγ, a key proinflammatory cytokine known to help control Mtb and lyse Mtb infected macrophages^38-41^. Due to their ability to home to pulmonary spaces and anti-Mtb effector functions, it is unsurprising that in preclinical models of Mtb challenge, CD8+ T cells help reduce bacterial burden^25,28,32,42^. Most recently, the use of intravenous (i.v.) BCG vaccination in nonhuman primates (NHP) has provided a benchmark of immune responses that correlated with nearly sterilizing protection from Mtb challenge^43^. Notably, in the routine intradermal (i.d.) BCG vaccine NHP cohort there was a lack of proinflammatory CD8+ T cells and less protection when compared to i.v. BCG^43^. These data collectively suggest CD8+ T cells represent an underappreciated target for vaccine-induced efficacy endpoints.

Many strategies designed to induce robust anti-Mtb CD8+ T cells have been developed and are well-reviewed here^25,26,30^. Some examples include, recombinant BCG (VPM1002), chimpanzee adenoviral vectored strategies (ChAdOx185A), modified vaccinia virus Ankara (MVA85A), recombinant adenoviruses (Ad5Ag85A, AERAS-402), and Cytomegalovirus vector approaches (CMV-6Ag and MTBVAC) and are under clinical evaluation^20,25,26,44,45^. Enriching the pipeline with novel candidates and platforms is warranted given the expanding immune correlates being discovered and need for broad accessibility in LMICs. Leveraging RNA vaccines for example, circumvents the challenges of protein manufacturing, reducing vaccine development timelines from several years to months and reducing costs throughout pipeline evaluations and deployment. For example, once the SARS-CoV-2 sequence was made available publicly, messenger RNA (mRNA) vaccines were developed within months and tested in human clinical trials^46^. While RNA vaccine platforms have been very minimally leveraged for TB vaccines^47^ to our knowledge, the evidence for targeting a multifaceted immune responses against Mtb for high vaccine efficacy provides good rationale for evaluating this platform. We and others have previously demonstrated robust humoral and cellular responses, including CD8+ T cell responses, induced by replicating RNA delivered via multiple different modalities^48-51^. For these reasons, we designed and evaluated a novel replicating RNA platform (repRNA)^48^ both as a homologous TB vaccine and as a heterologous strategy to complement protein – adjuvant prophylactic vaccine candidates against Mtb challenge in mice.

## Results

### Mtb RNA Candidate Vaccine

Preclinical Mtb vaccine candidate, ID91, is a fusion of four Mtb proteins: Rv3619 (esxV; ESAT6-like protein), Rv2389 (RpfD), Rv3478 (PPE60) and Rv1886 (Ag85B) (**Figure 1A**). These antigens were selected as candidates for Mtb vaccines based on their lack of human sequence homology and their ability to induce an *ex vivo* IFN-γ response in PPD+ human peripheral blood mononuclear cells (PBMC), suggesting they are immunogenic in humans^52,53^. ID91 antigens were cloned into an alphavirus repRNA backbone as previously described^48^ (**Figure 1B**). This repRNA platform has a number of advantages including 1) a subgenomic promoter to amplify expressioin of the antigen of interest, 2) exclusive cytoplasmic activity with no risks of nuclear genomic integration, and 3) sustained antigen production in the presence of adjuvanting innate responses that allows for dose sparing and strong cellular responses^48,49^. Using western blotting we observed efficient expression of the ID91 antigens from repRNA ID91 in BHK cells (**Figure 1C**), demonstrating this platform is amenable to Mtb antigen expression. Partnering repRNA with delivery formulations provides stability and protection from RNAses and removes the requirement for viral delivery, eliminating anti-vector immunity which has been detrimental for promising candidates. Here we leveraged a first-generation nanostructured lipid carrier (NLC)^48^ as a two-vial approach with ID91-repRNA. C57BL/6 mice were vaccinated two times, two weeks apart with either ID91 fusion protein + TLR4 agonist, glucopyranosyl lipid adjuvant formulated in a stable emulsion (GLA-SE) or ID91-repRNA + NLC to evaluate preliminary immunogenicity. Splenocytes from both immunized groups were stimulated two weeks post boost with media alone, ID91 protein or each of 13 different 15mer peptide pools overlapping by 8mers of ID91 and evaluated for proliferation and cytokine responses (**Figure 1D**). We observed that ID91+GLA-SE elicited both CD4+ (Red) and CD8+ (Blue) T cell responses, albeit larger CD4+ TH1, while ID91-repRNA+ NLC largely drove CD8+ T cell proliferation and inflammatory cytokines (**Figure 1D**). While there is some overlap in CD4+ and CD8+ responses driven by each immunization strategy, these data demonstrate that RNA and protein differentially elicit CD4+ and CD8+ T cellular responses to distinct antigen epitopes (**Figure 1D**) which may be beneficial for heterologous vaccine approaches. This is congruent with clinical trials of ID93+GLA-SE where we have observed protein-adjuvant vaccine-elicited immunogenicity to be dominated by CD4+ TH1 responses and far fewer cases (4 responders of 38 across all regimens tested) demonstrating CD8+ T cell based responses^54^.

**Figure 1.**
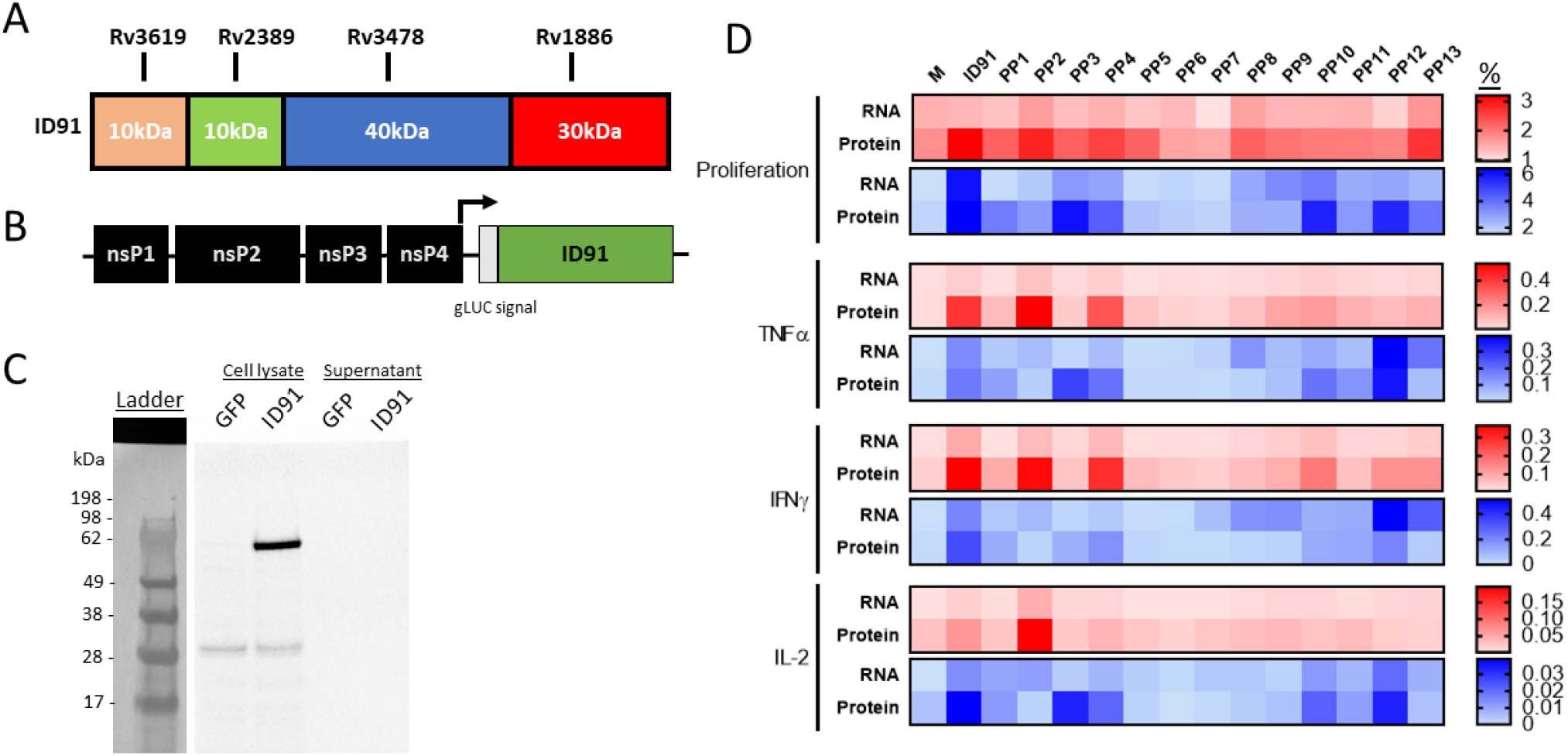
Design and Characterization of ID91 repRNA vaccine candidate. **A**) ID91 fusion protein comprising 4 Mtb Antigens Rv3619, Rv2389, Rv3478, Rv1886. **B**) Alphavirus backbone expressing nonstructural proteins (nsP) and ID91 fusion antigen. **C**) Western blot of ID91 antigen expression from transfected BHK cell lysate and supernatant *in vitro* after 24 hours. **D**) Epitope mapping of splenocytes from ID91 repRNA + NLC (RNA) or ID91+GLA-SE immunized C57BL/6 mice. Splenocytes were stimulated with media (M) whole antigen (ID91) or 13 different peptide pools (PP) with overlapping 15mers of ID91. Proliferation and cytokine expression were measured by flow cytometry after stimulation. Heat map depicts percentage of CD4+ (red) or CD8+ (blue) responding T cells from each immunization depicted to the left. Data representative of a single experiment.

### Immunogenicity and Prophylactic Protective Efficacy

We first evaluated ID91 repRNA and ID91-GLA-SE prophylactic immunizations in C57BL/6 mice for immunogenicity as well as protection from infection six weeks after a single immunization as a stringent criteria for advancing this platform. A single vaccination with ID91+GLA-SE induced robust CD4+ CD44+ TH1 T cells expressing IFNγ, IL-2 and TNF (**Figure 2A**), but negligible CD8+ CD44+ cytokine producing T cells (**Figure 2B**). Conversely, ID91 repRNA + NLC trended towards higher induction of IFNγ, IL-2 or TNF producing CD8+ CD44+ T cells (**Figure 2B**) as well as moderate generation of TH1 CD4+ T cell responses (**Figure 2A**). No significant IL17a or IL-21 expression was observed for any group (data not shown). Both vaccines induced a robust total IgG humoral immune response to the ID91 fusion, significantly greater than the saline group but not significantly different between the two platforms, with the bulk of this response being against antigen Rv1886 (**Figure 2C**). Lastly, we observed that both ID91+GLA-SE and ID91 repRNA + NLC significantly reduced lung bacterial burden, 0.55 and 0.43 log protection versus saline respectively, three weeks post challenge with a low-dose aerosol (LDA 50-100 CFU) of Mtb H37Rv (**Figure 2D**). These data demonstrate that the repRNA platform is immunogenic and affords some prophylactic protection in a stringent preclinical mouse efficacy screening model of TB.

**Figure 2:**
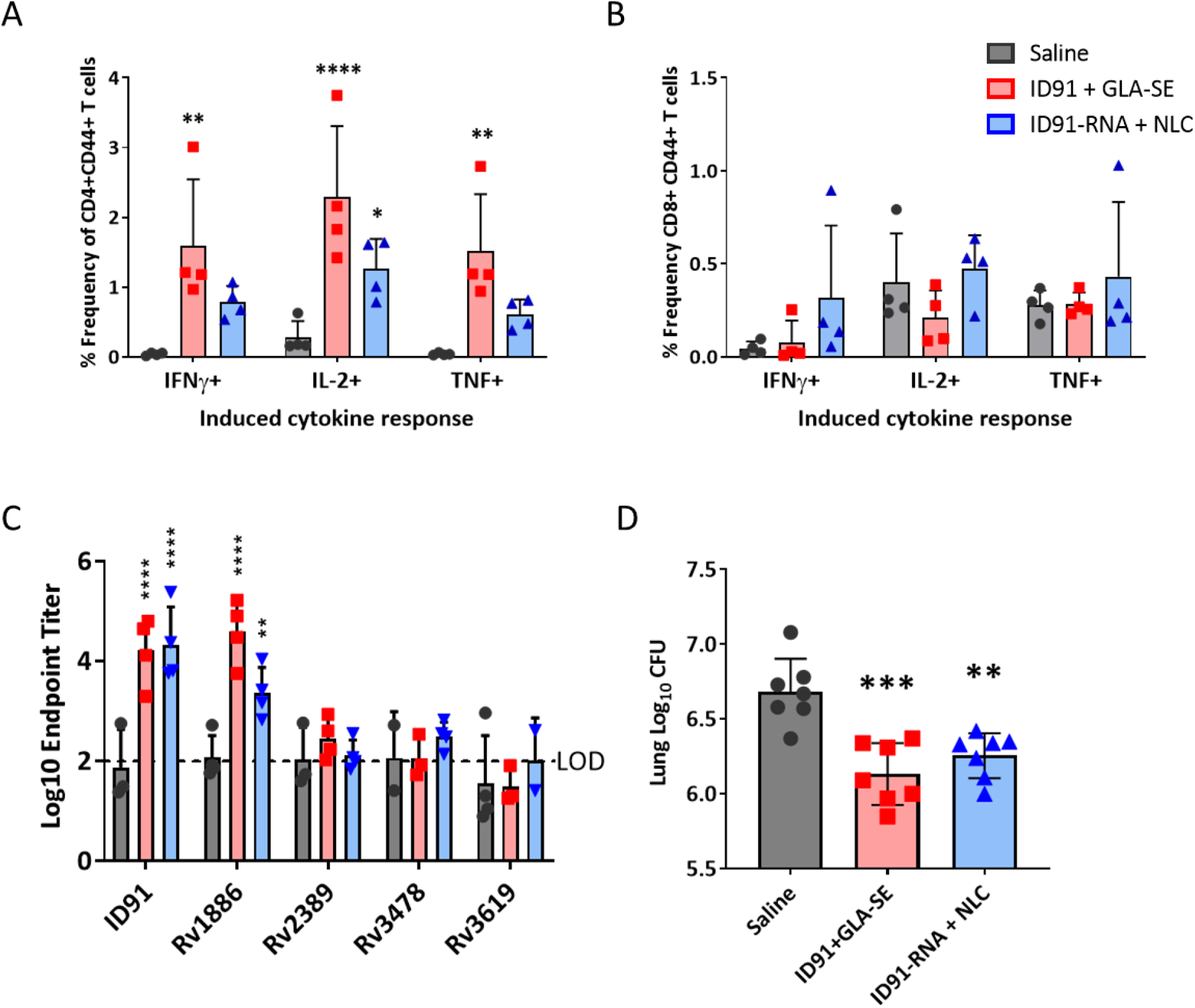
Single immunization with repRNA vaccine is moderately immunogenic and affords prophylactic protection. Splenocytes from cohorts of saline (grey circles), ID91+GLA-SE (red squares) or ID91 repRNA + NLC (blue triangles) immunized mice 6 weeks post-vaccination were stimulated with ID91 *ex vivo* and evaluated by intracellular cytokine staining flow cytometry including **A**) percentage of CD4+ CD44+ cytokine producing T cells as well as **B**) percentage of CD8+ CD44+ cytokine producing T cells. n=4 per group. **C**) Plasma was evaluated 6 weeks post vaccination for total IgG responses to fusion antigen ID91 and its components, Log10 EPT is shown. n=4 per group. **D**) Pulmonary bacterial burden Log10 CFU 3 weeks post LDA challenge with Mtb H37Rv. n=7 per group. Asterisks represent a statistically significant difference from saline using One-way ANOVA with Dunnett’s multiple-comparison test. All data are representative of two independent repeated experiments. * p < 0.05, ** p < 0.01, *** p < 0.001 and **** p < 0.0001.

Next, we evaluated the protein and RNA-based platforms for immunogenicity and protection against an ultra-low dose (ULDA, 1-8 bacteria) of H37Rv challenge using heterologous, homologous and combination prime-boost strategies. For combination strategies mice were given both ID91+GLA-SE and ID91 repRNA + NLC i.m. in separate limbs at prime and boost time points to avoid potential innate immune-mediated interference between the two modalities. We hypothesized that driving CD4+ and CD8+ T cell responses may have an additive or synergistic effect on immunogenicity and subsequent protection from the more physiologically relevant ULDA challenge. We observed that two immunizations with ID91+GLA-SE enhanced protection in the lung (0.743 log10) over a single dose (**Table 1, Figure 3A**) which is in line with prior publications from our group^55^. However, two immunizations with ID91 repRNA + NLC only afforded moderate reduction of bacterial burden in the lung, 0.306 log10 (**Table 1, Figure 3A**), which we believe may be due to suboptimal timing between prime and boost vaccinations based on recent observations outside the scope of this manuscript. Heterologous strategies also differed, with RNA-prime protein-boost being the most efficacious regimen evaluated for reduction of lung bacterial load (0.847 log10 reduction), followed closely by the combination regimen (0.809 log10 reduction) (**Table 1, Figure 3A**). While the combination regimen appears most protective from dissemination to the spleen (1.419 log10 reduction from saline), this was not statistically significant due to a wide standard error (**Table 1, Figure 3B**).

**Figure 3.**
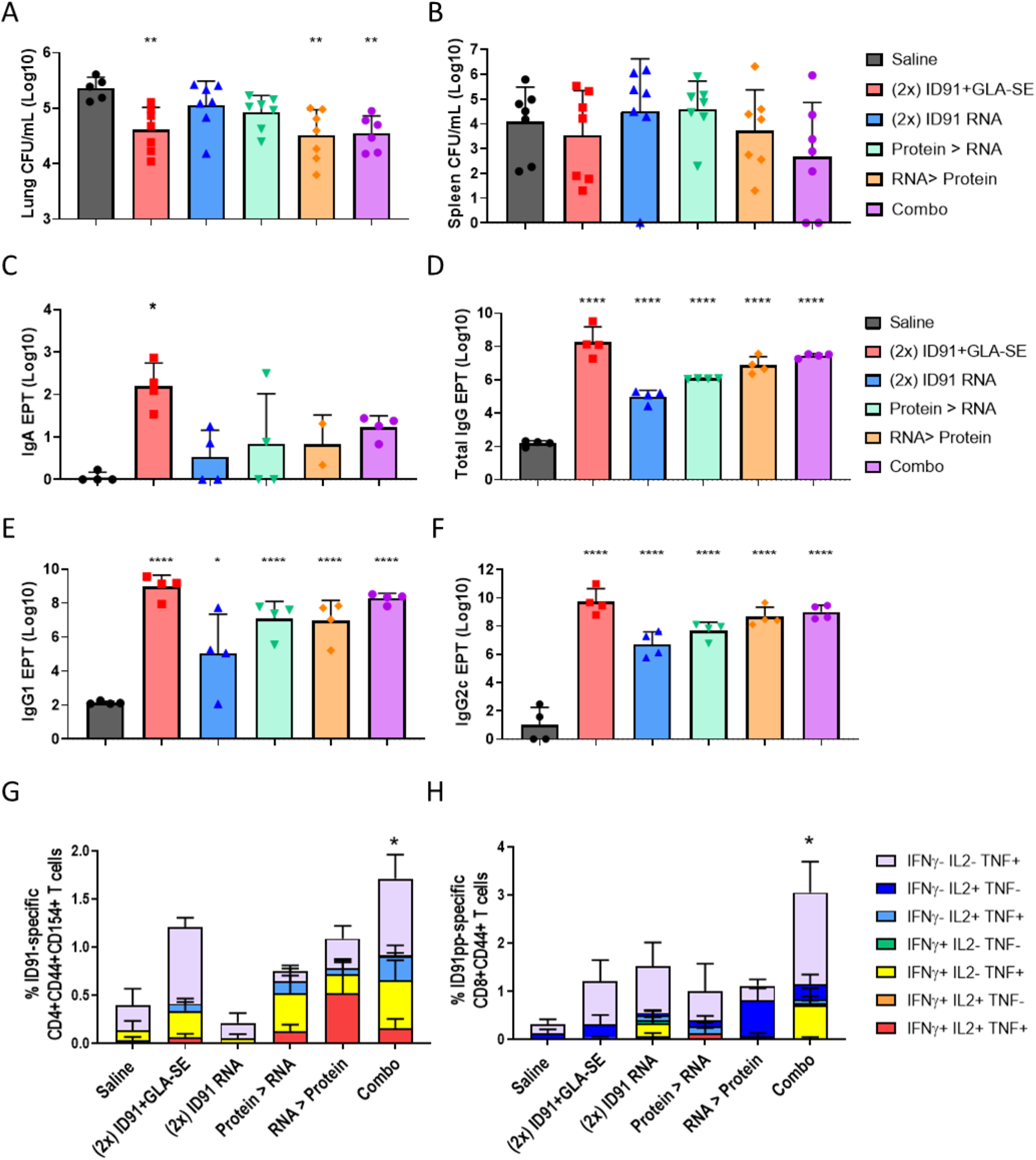
Regimen composition influences protection and post challenge immune responses. Bacterial burden assessed by colony forming unity (CFU) in **A)** lung and **B)** spleen homogenates 3 weeks post challenge. n=6-7 mice per cohort, saline – grey, homologous ID91+GLA-SE – red, homologous ID91 repRNA +NLC – blue, protein prime/RNA boost – green, RNA prime/protein boost – orange, and combination – purple. **C)** BALf from n=4 mice per cohort was collected 3 weeks post challenge and evaluated for ID91-specific IgA responses. Plasma from n=4 mice per cohort was isolated 3 weeks post challenge and evaluated for ID91-specific **D)** Total IgG, **E)** IgG1 and **F**) IgG2c. CFU and EPT group means were compared to saline by ordinary One-Way ANOVA and Dunnet’s multiple comparisons test. Single cell suspensions isolated from the lungs of animals 3 weeks post ULDA challenge were stimulated with ID91 protein (CD4+ T cells) or ID91 peptide pool (CD8+ T cells) and evaluated for **G)** CD4+CD44+CD154+ and **H)** CD8+CD44+ T cell expression of IFNγ, IL-2 and TNF by flow cytometry. n=4 per group. All data are representative of two independent repeated experiments. Cytokine expression was compared using a One-Way ANOVA and Dunnet’s multiple comparisons test. * p < 0.05, ** p < 0.01, *** p < 0.001 and **** p < 0.0001.

**Table 1.**
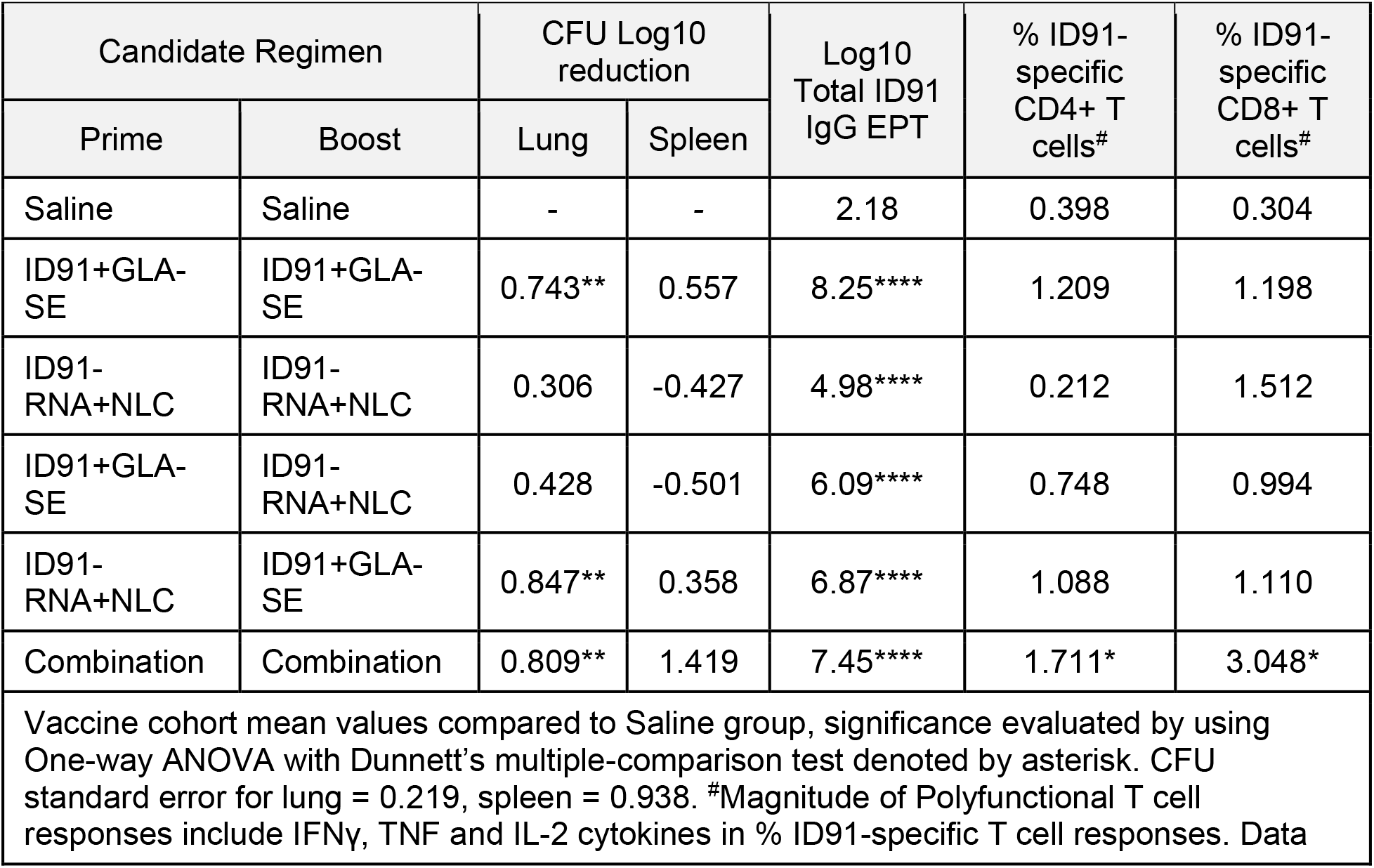
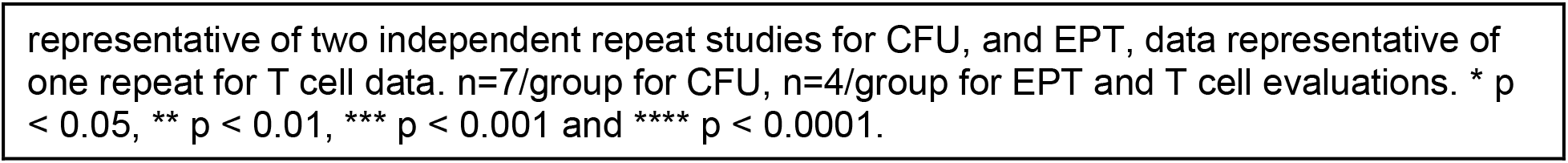
Immunogenicity and Efficacy using Prime-Boost Regimens.

This pattern of efficacy is largely mirrored in the post-challenge humoral and cellular immunogenicity readouts. While i.m. vaccine strategies are not known for developing robust mucosal immune responses, we evaluated bronchoalveolar lavage fluid for anti-ID93 IgA responses at 3 weeks post challenge. Not surprisingly the magnitude of IgA produced by all regimens were low, and homologous ID93+GLA-SE was the only strategy to induce a statistically significant response compared to saline (**Figure 3C**). We observed that combination and homologous ID91+GLA-SE regimens induced the highest ID91-specific total IgG endpoint titers in serum (EPT) post challenge, 7.45 log10 and 8.25 log10 respectively (**Table 1, Figure 3D**). They are followed by heterologous regimens, while the lowest total IgG induced was in the homologous RNA cohort (**Table 1, Figure 3D**). Similar trends were observed for ID91-specific IgG1 (**Figure 3E**) and IgG2c (**Figure 3F**), with homologous protein and combination regimens demonstrating the highest EPT and homologous RNA the lowest EPT.

Interestingly we did not observe significant differences in ID91-specific stimulation of individual cytokines IL-2, GM-CSF or IL-17A from CD4+ nor CD8+ T cells between treatment groups. However, CD4+ T cell TNF expression was statistically higher in heterologous RNA prime, protein boost and combination regimens and CD8+ T cell TNF expression was significantly higher in the combination regimen. Given the relative paucity of strong CD8+ epitopes in ID91 antigens, it was not a surprise to observe expression of CD107a on CD8+CD44+ T cells post challenge to be statistically similar between cohorts (data not shown). We also examined polyfunctional CD4+ and CD8+ T cell responses by flow cytometry from each cohort after *ex vivo* stimulation of lung cells and found only the combination regimen to induce significantly higher total magnitude of responses for both CD4+ and CD8+ subsets (**Figure 3G, 3H, Table 1**). While not significant, there is a trend for most robust polyfunctional CD4+ T cell responses in regimens that include at least one arm including protein immunizations. We observed the composition of polyfunctional cells to be relatively similar (**Figure 3G, 3H**). Triple positive (IFNγ, IL-2 and TNF), dual expressing IFNγ+ TNF+ and single TNF positive cells made up the majority of the CD4+ T cell responses across regimens evaluated, while single positive IL-2+ or TNF+ responses overwhelmingly dominated the CD8+ T cell response in this post challenge *ex vivo* stimulation (**Figure 3G, 3H**).

## Discussion

These data provide the first report of a replicating RNA-based vaccine strategy being evaluated in the Mtb mouse challenge model and showing efficacy in heterologous and combination regimens. A key advantage of RNA platforms is their ability stimulate antigen-specific CD8+ T cells and antibody responses which both aid in the control of intracellular infections. As a result, heterologous and co-immunization strategies that combine protein and nucleic acid vaccines is a promising approach to inducing protective immunity through CD4+ and CD8+ T cell-mediated responses^56,57^. We observed that heterologous and combination regimens were moderately additive in reducing pulmonary bacterial burden in this preclinical model. While no single immunological endpoint (total IgG, IgG1, IgG2c, IgA EPT, TH1 CD4+ or CD8+ T cells) correlated with protective efficacy, there are interesting trends that may be further explored and optimized. Indeed, our data suggest that meeting thresholds for specific combinations of responses, versus single endpoints, may better correlate with protective efficacy in this model. Additionally, future interrogations should examine other immune correlates that are showing promise including IgM, IgA as well as localization and kinetics of CD4+ and CD8+ T cell responses^43,58^. The mucosal or aerosol delivery of repRNA in specialized delivery formulations may also help drive pulmonary-specific immunity against Mtb and these strategies are currently being evaluated.

The landscape of TB vaccines is ready for a dramatic surge forward. While ID91 served here as a proof-of-concept antigen which afforded some protection from bacterial burden, epitope mapping demonstrated that ID91 contains few dominant CD8+ T cell epitopes in the C57BL/6 mouse model. Future work that incorporates antigens containing high priority CD8+ T cell epitopes^29,59,60^ should be prioritized. The speed of RNA vaccine development and ability to rapidly swap antigens lends itself well to the need of TB vaccines since global regions harbor different predominant lineages^61^ and tailoring vaccines to meet regional needs may further help improve efficacy. Additionally, evaluating alternative repRNA expression strategies and alternative timing between prime and boost immunizations could be explored to optimize B and T cell responses to the delivered antigens. For example, here we evaluated a prime/boost spaced 3 weeks apart and have more recently established that 4-8 weeks between prime and boost drives better immune responses in regimens that include the repRNA platform (unpublished and ^49^). We believe that the moderate additive effect seen here with heterologous and combination regimens could be significantly improved with these strategies. Furthermore, while this preclinical proof of concept work leveraged an NLC formulation, next generation RNA vaccine formulations have advanced into clinical trials (e.g. HDT BioCorp: state-of-the-art delivery vehicle Lipid InOrganic Nanoparticle (LION)^49^, US ClinicalTrials.gov Identifier: NCT04844268, India CTRI/2021/04/032688) and development of RNA-based Mtb vaccines should leverage these formulations or others^62-66^ with proven human safety data to help accelerate this platform to address the ongoing TB epidemic. In summary, the repRNA platform shows promise as a vaccine strategy for TB and a well-designed set of Mtb antigens in partnership with advanced RNA formulations have the capability to significantly contribute to the TB vaccine pipeline.

## Materials and Methods

### ID91 repRNA production and qualification

The codon optimized sequence encoding ID91 was synthesized (Codex DNA) and cloned into the Venezuelan equine encephalitis virus replicon (strain TC83) between PflFI and SacII sites by Gibson assembly. Sequence-verified DNA was then linearized by NotI digestion and RNA transcribed and capped by T7 RNA polymerase and Vaccinia capping enzyme reactions, respectively, as previously described^49^. To validate the ID91 repRNA produced ID91 protein, Baby Hamster Kidney (BHK) cells (ATCC) were transfected with 4 µg ID91-repRNA or GFP-repRNA as a negative control using lipofectamine 2000 (ThermoFisher). After 24 hours cells and supernatants were collected for analysis by SDS-PAGE and Western blot under reducing and non-reducing conditions. For ID91 antigen detection after transfer to PVDF membrane, primary mouse sera (from animals immunized 3 times 3 weeks apart with ID91+GLA-SE) was used at 1:500 in PBS. Goat anti-mouse HRP (SouthernBiotech) 1:10000 was used for detection along with SuperSignal™ West Pico PLUS Chemiluminescent Substrate (ThermoFisher). The resulting image was captured on a Biorad chemidoc XRS+.

### Preclinical animal model

Female C57BL/6 mice 4-6 weeks of age were purchased from Charles River Laboratory. Mice were housed at the Infectious Disease Research Institute (IDRI) biosafety level 3 animal facility under pathogen-free conditions and were handled in accordance with the specific guidelines of IDRIs Institutional Animal Care and Use Committee. Mice were infected either with a low dose (50–100 bacteria) aerosol (LDA) or ultra-low dose (1-8 bacteria) aerosol (ULDA) of Mtb H37Rv using a University of Wisconsin-Madison aerosol chamber. Twenty-four hours post challenge the lungs of 3 mice were homogenized and plated on Middlebrook 7H11 agar (Fisher Scientific) to confirm delivery.

### Vaccines and adjuvants

Cohorts of mice were immunized intramuscularly (i.m.) twice three weeks apart for epitope mapping, once with 6 weeks before challenge, or twice three weeks apart, with the final immunization occurring 3 weeks before challenge for immunogenicity and efficacy testing. Mice received either saline alone, or vaccinations containing 0.5 µg/dose of ID91 recombinant fusion protein combined with 5.0 µg/dose GLA-SE, as previously published^67-69^. A separate cohort of mice were immunized with 1.0 µg repRNA complexed with NLC as described^48^. Homologous or heterologous regimens were also leveraged using the doses and regimens outlined above. Combination cohorts received ID91+GLA-SE in the right hind limb followed by ID91 repRNA + NLC in the left hind limb at both prime and boost timepoints in the same doses described above.

### Epitope mapping

Two weeks post boost splenocytes were isolated from cohorts of ID91+GLA-SE or ID91 repRNA + NLC immunized mice and stimulated with ID91 protein antigen (10µg/mL), overlapping peptides (10µg/mL) or media alone for 60 hours at 37 °C and 5% CO_2_ with proliferation dye (Violet Proliferation Dye, BD Biosciences). Peptides were generated in 15mer format with 8 amino acid overlaps for the ID91 fusion antigen, for a total of 125 peptides (BioSynthesis). These 125 peptides make up 13 different pools of 5-10 peptides each. Cells were then restimulated for two hours, brefeldin A was added directly to the plate and cells were allowed to incubate for a further 4 hours. Cells were then washed and ICS and flow cytometry analysis were performed as described below.

### Flow cytometry

Intracellular flow cytometry was performed on splenocytes post immunization but pre-challenge and lung homogenates post infection. Samples were incubated, washed and stimulated as previously described^68^. Cells were stimulated with media alone, 10 µg/mL of recombinant ID91, or 1 µg/mL phorbol myristate acetate (PMA) (Calbiochem) + 1 µg/mL ionomycin (Sigma-Aldrich). Some sample stimulations also contained fluorescently labeled anti-mouse CD107a (1D4B, BioLegend). After stimulation samples were stained for markers of interest using fluorescent conjugated antibodies as previously described^68^. Notably, all sample stains were used at 10 µL/mL concentration. Primary surface staining included: anti-mouse CD4 (clone RM4-5, BioLegend), CD8a (clone 53-6.7, BioLegend), CD44 (clone IM7, eBiosciences), CD154 (clone MR1, BioLegend) and 1 µg/mL of Fc receptor block anti-CD16/CD32 (clone 93, eBioscience) in PBS with 1% bovine serum albumin (BSA). Cells were then washed and fixed using BD Biosciences Fix/Perm reagent for 20 min at RT. Intracellular staining was carried out in Perm/Wash (BD Biosciences) reagent with anti-mouse GMCSF (clone MP1-22E9, BioLegend), IFN-γ (clone XMG1.2, Invitrogen), IL-2 (clone JES6-5H4, eBioscience), IL-17A (clone Tc11-1BH10.1, BioLegend), TNF-α (clone MP6-XT22, eBioscience), and IL-21 (mhalx21, eBiosciences) for 10 min at RT. Samples were then incubated in 4% paraformaldehyde for 20 min to fix, decontaminate and remove from the containment facility before washing and resuspension in PBS + 1% BSA. An LSRFortessa flow cytometer (BD Bioscience) was used for sample acquisition, analysis was performed using FlowJo v10.

### EPT ELISA

Serum was collected from terminal bleeds immediately after euthanasia, at post boost or post infection timepoints. BALf was collected from post infection timepoints immediately after euthanasia. Serum samples were diluted 1:100, and BALf samples were added directly (neat) to plates coated with 2 µg/mL of ID91, Rv1886, Rv2389, Rv3478, or Rv3619. After an overnight incubation, plates were exposed to HRP-conjugated IgA, Total IgG, IgG1, or IgG2c (Southern Biotech). Plates were then developed with tetramethylbenzidine substrate and stopped with 1 N H_2_SO_4_. Plates were read at 450nm with 570nm background subtraction. EPT were calculated using regression analysis of sample dilution and O.D. on GraphPad Prism.

### Bacterial burden/CFU

24 hours to 3 weeks post infection with Mtb, 3–7 mice per group were euthanized with CO_2_. Lung and spleen tissue from infected animals was isolated and homogenized in 5 mL of either RPMI + FBS (lung) or PBS + Tween-80 (Sigma-Aldrich, St. Louis, Missouri, USA) CFU buffer (spleen) using an Omni tissue homogenizer (Omni International, Kennesaw, GA, USA). Serial dilutions of homogenate were made in CFU buffer and aliquots were plated on Middlebrook 7H11 agar plates and subsequently incubated at 37 °C and 5% CO_2_ for 2– 3 weeks before colonies were counted. Bacterial burden, as CFU/mL, was calculated per organ and is presented here as Log10 values. Reduction in the bacterial burden was calculated as the difference in mean Log10 values between groups assessed.

### Statistics

Bacterial burden, humoral and cellular immune responses were assessed using a One-way ANOVA with Dunnets multiple comparison test between vaccinated groups and saline control. Flow cytometry data was assessed using FlowJo v10 (BD) and SPICE (NIH) using the Wilcoxon signed rank test as compared to untreated groups or One-way ANOVA with Dunnets multiple comparison test. Statistical analyses were performed using GraphPad Prism 7 software. Significant differences are labeled accordingly in the figures where * p < 0.05, ** p < 0.01, *** p < 0.001 and **** p < 0.0001, with methodology used previously by our group^68,70^.

## Acknowledgements

The authors would like to express their gratitude to Seattle Children’s Research Institute Leadership for their guidance, compassion, and mentorship.

The datasets generated and/or analyzed during the study presented here are available from the corresponding authors on reasonable request.

Research reported here was supported by the National Institute of Allergy and Infectious Diseases (NIAID) of the National Institutes of Health (NIH) under award number R01AI125160. SEL was supported through the University of Washington, Diseases of Public Health Importance T32 training grant number A1075090. The content is solely the responsibility of the authors and does not necessarily represent the official views of the National Institutes of Health.

## Author Contribution Statement

SEL, JHE, SLB and RNC designed experiments; SEL, JHE, VAR, TP, JA, AK, FCH carried out experiments; SEL, VAR and TP executed statistical analysis; SEL drafted the manuscript; SEL, JHE, VAR, TP, JA, FCH prepared figures and tables. All authors (SEL, JHE, VAR, TP, JA, AK, FCH, SGR, SLB, RNC) reviewed the manuscript and contributed to edits.

## Additional Information

All authors declare no financial or alternative competing interests with the data described in this manuscript.

